# Optimizing bulk segregant analysis of drug resistance using *Plasmodium falciparum* genetic crosses conducted in humanized mice

**DOI:** 10.1101/2021.11.24.469878

**Authors:** Katelyn V. Brenneman, Xue Li, Sudhir Kumar, Elizabeth Delgado, Lisa A. Checkley, Douglas A. Shoue, Ann Reyes, Biley A. Abatiyow, Meseret T. Haile, Rupam Tripura, Tom Peto, Dysoley Lek, Katrina A Button-Simons, Stefan H. Kappe, Mehul Dhorda, François Nosten, Standwell C. Nkhoma, Ian H. Cheeseman, Ashley M. Vaughan, Michael T. Ferdig, Tim J. C. Anderson

## Abstract

**Background:** Classical genetic crosses in malaria parasites involve isolation, genotyping, and phenotyping of multiple progeny parasites, which is time consuming and laborious. Bulk segregant analysis (BSA) offers a powerful and efficient alternative to identify loci underlying complex traits in the human malaria parasite, *Plasmodium falciparum.*

**Methods:** We have used BSA, which combines genetic crosses using humanized mice with pooled sequencing of progeny populations to measure changes in allele frequency following selection with antimalarial drugs. We used dihydroartemisinin (DHA) drug selection in two genetic crosses (*Mal31×KH004* and *NF54×NHP1337*). We specifically investigated how synchronization, cryopreservation, and the drug selection regimen of progeny pools impacted the success of BSA experiments.

**Findings:** We detected a strong and repeatable quantitative trait locus (QTL) at chr13 *kelch13* locus in both crosses, but did not detect QTLs at ferredoxin (*fd*), the apicoplast ribosomal protein S10 (*arps10*), multidrug resistance protein 2 (*mdr2*). QTLs were detected using synchronized, but not unsynchronized pools, consistent with the stage-specific action of DHA. We also successfully applied BSA to cryopreserved progeny pools.

**Interpretation:** Our results provide proof-of-principal of the utility of BSA for rapid, robust genetic mapping of drug resistance loci. Use of cryopreserved progeny pools expands the utility of BSA because we can conduct experiments using archived progeny pools from previous genetic crosses. BSA provides a powerful approach that complements traditional QTL methods for investigating the genetic architecture of resistance to antimalarials, and to reveal new or accessory loci contributing to artemisinin resistance.

**Funding:** National Institutes of Health (NIH); Wellcome trust.

Research in context

Evidence before this study
Genetic crosses have been immensely successful for determining the genetic basis of drug resistance in malaria parasites, but require laborious cloning, characterization of drug resistance and genome-wide genotyping of individual progeny. This is a major limitation given that genetic crosses can now be conducted efficiently using humanized mice, rather than chimpanzees. Bulk segregant analysis (BSA) provides an attractive alternative approach because (i) large numbers of uncloned recombinant progeny can be analyzed, increasing statistical power (ii) phenotyping is not required, because we identify QTLs by treatment of progeny bulks, and identifying genome regions that show skews in allele frequency after treatment (iii) genome sequencing of bulk samples provides a rapid, accurate readout of genome-wide allele frequencies. This approach has been effectively leveraged in yeast, rodent malaria and several model organisms.

Added value of this study
Here we validate and optimize this approach for *P. falciparum* genetic crosses, focusing on resistance to dihydroartemisin (DHA) a central component of the first-line antimalarial combination. Mutations in *kelch13* are known to confer resistance to DHA, but several additional candidate loci have also been suggested to contribute. Our results confirm involvement of *kelch13*, but did not identify linkage with other putative candidate loci. We optimized methodology, showing that synchronization is critical, and that BSA can be successfully applied to cryopreserved progeny pools.

Implications of all the available evidence
BSA combined with recent advances in rapidly generating genetic crosses provides a powerful approach to investigate the genetic basis of drug resistance in *P. falciparum*.

## 1. Introduction

Identifying genetic changes conferring drug resistance helps understand the resistance mechanisms and track the spread of resistance alleles in infectious disease-causing organisms, including malaria parasites. For malaria parasites, both association mapping and linkage analysis approaches have been used to understand the genetic mechanisms underlying drug resistance ^1^. For example, the chloroquine resistance transporter (*crt*) conferring chloroquine (CQ) resistance in the malaria parasite *Plasmodium falciparum* was initially located on chromosome 7 through linkage analysis ^2,3^ and further identified by fine mapping ^4^. Genome-wide association studies (GWAS) have also been applied to map genes associated with resistance to antimalarials ^5^ as well as additional loci arising on genetic backgrounds that carry resistance-associated alleles ^6^. However, both linkage mapping and association studies have their limitations: linkage mapping requires laborious isolation, genotyping, and phenotyping of multiple progeny parasites; GWAS requires months or years of sample collection.

Bulk segregant analysis (BSA), a complementary alternative to traditional linkage methods, can provide a simple and rapid approach to identify loci that contribute to complex traits without the need to phenotype and genotype individual progeny. Instead, using pooled sequencing of progeny populations, BSA measures changes in allele frequency following the application of different selection pressures (e.g., by drugs). The BSA approach is simple; by using the complete pool of unique recombinants for analysis it requires less labor and cost and has the potential for increased statistical power over traditional linkage mapping ^7^. Nevertheless, BSA cannot directly examine epistatic interactions or measure phenotypes that are not amenable to selection in bulk. The BSA approach was initially developed to study diseases in human and plants, in which DNA from individuals with different phenotypes were pooled and genotyped to identify loci that were enriched for different alleles in each pool ^8^. BSA has also been extensively used with genetic crosses of rodent malaria parasites (where it is termed linkage group selection (LGS)) to map genes determining blood stage multiplication rate, virulence, and immunity in *Plasmodium yoelii* ^9^, as well as mutations conferring ART resistance and strain-specific immunity in *Plasmodium chabaudi* ^10,11^. BSA has also been used in yeast ^7^, *C. elegans* ^12^, *Eimeria tenella* ^13^, and the human blood fluke *Schistosoma mansoni* ^14^. We have previously applied BSA to the human malaria parasite *P. falciparum* to identify genes that impact parasite fitness throughout the intra-erythrocytic stages of the lifecycle ^15^ and polymorphisms in nutrient acquisition/metabolism pathways ^16^.

Artemisinin combination therapies (ACTs) are the current first-line malaria treatments in most countries for the most lethal human malaria parasite, *P. falciparum.* Resistance to artemisinin (ART) is mediated by mutations in *kelch13,* a gene required for parasite endocytosis ^17^. Studies in Southeast Asia uncovered the independent emergence of *kelch13* resistance-associated alleles and their rapid spread ^18, 19^ Furthermore, recent studies have also detected an increased allele frequency of *kelch13* mutations in clinically artemisinin-resistant *P. falciparum* in Africa ^20^. Nonsynonymous polymorphisms in ferredoxin (*fd*), the apicoplast ribosomal protein S10 (*arps10*), multidrug resistance protein 2 (*mdr2*), and *crt* have also been identified and show strong associations with artemisinin resistance ^6^, while (iv) *in vitro* selection has revealed other loci, such as *pfcoronin*, that can modulate ART-R ^21^.

We designed this study to optimize aspects of the BSA approach for the identification of drug resistance-associated loci in malaria parasites (**Figure 1**), using dihydroartemisinin (DHA), the active metabolite of artemisinin, as our test drug. We chose DHA because (i) artemisinin is one of the most widely used antimalarial drugs, (ii) *kelch13* is a known resistance locus, providing a good control for our methods, (iii) identification of additional loci involved in drug resistance would be of great interest, and (iv) this drug is stage specific, targeting early rings, providing additional challenges for BSA. We aimed to answer three main questions. First, for stage-specific drugs such as DHA, can we detect QTLs in unsynchronized parasite pools, or is prior synchronization required? This is important because synchronization may reduce the size of progeny pools, and for successful BSA it is critical to have progeny pools containing large numbers of recombinants. Second, how can we determine the optimal drug dose regimen to maximize signal? Third, can we use cryopreserved progeny pools for BSA experiments? This would greatly enhance the utility of BSA by allowing experiments to be conducted long after genetic crosses have been conducted, using archived progeny pools. However, as with synchronization, we were concerned that cryopreservation would reduce the diversity of progeny pools limiting our ability to detect QTLs.

**Figure 1.**
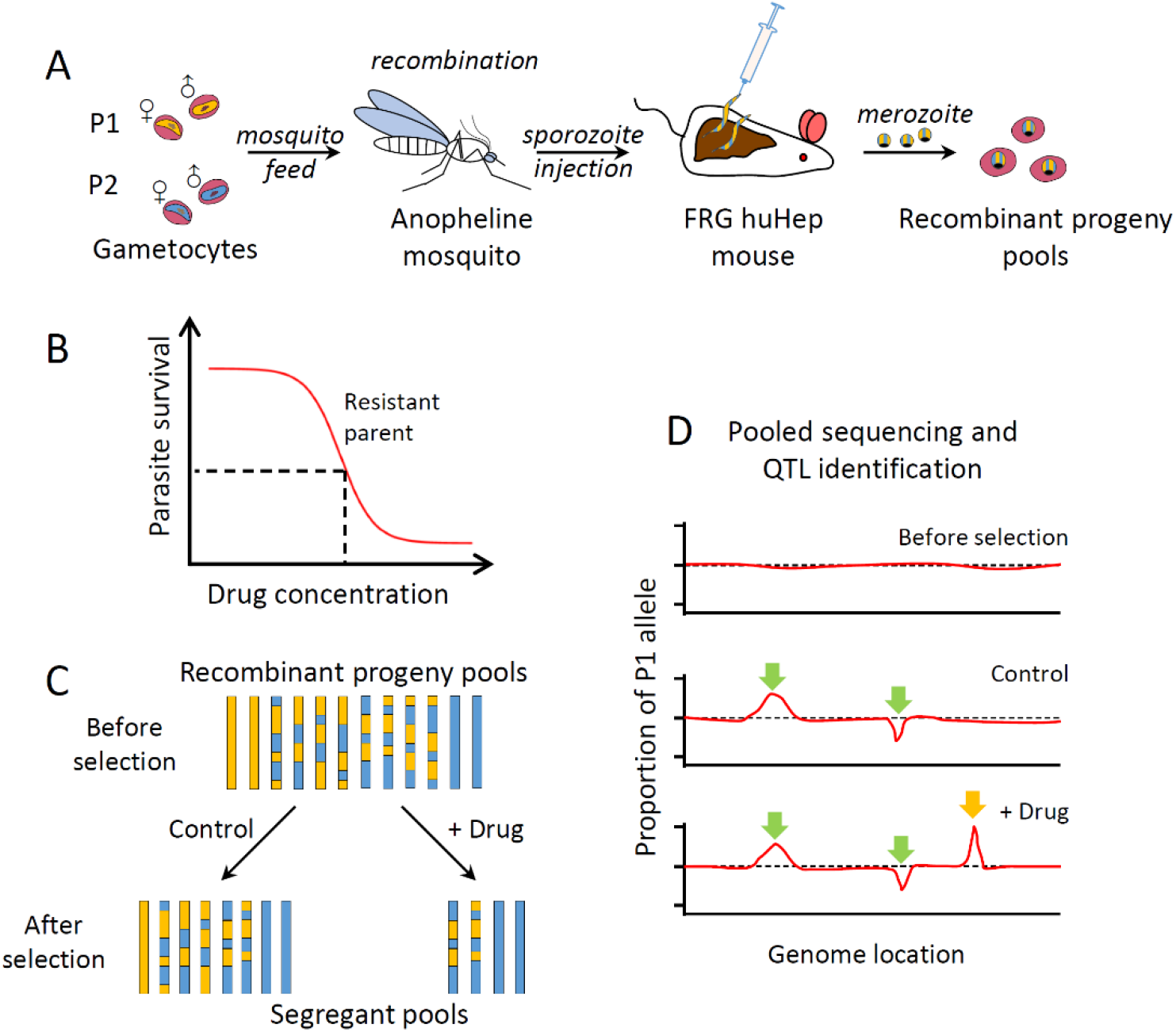
The principles of bulk segregant analysis with human malaria parasites. (A) Generation of recombinant progeny pools using *Anopheles stephensi* mosquitoes and FRG huHep mice. (B) Determination of drug concentration for bulk segregant analysis (BSA). A standard approach is to measure the IC_50_ and IC_90_ of the resistant parent, however, in this study, we used eRRSA to determine a discerning drug concentration for selection since classical IC_50_ and IC_90_ measurements using DHA do not correlate with clinical parasite clearance rate. (C) Apply selection with drug of choice, each bar indicates a representative recombinant progeny. (D) Genomic sequencing measures changes in allele frequency of segregant pools. Green arrows indicate allele frequency changes detected from both pools with and without drug treatment, which are thus not related to drug resistance, while the orange arrow indicates a locus conferring drug resistance.

We performed DHA BSA in two genetic crosses between ART-S African and ART-R Southeast Asian parasites (*Mal31×KH004* and *NF54×NHP1337*). These two crosses are of particular interest because SNPs in the *kelch13* locus, as well as SNPs in each of the four loci associated with ART-R in Miotto et al ^6^ genome wide association study (WGAS) are segregating in both crosses. We detected strong and repeatable QTL at the *kelch13* locus on chromosome 13 in both crosses, but found no evidence for involvement of other loci. Chr 13 QTLs were detected only in synchronized parasites. We also show that the application of eRRSA, a modification of the ring stage survival assay, allows for a rational determination of optimal drug dose for use in BSA. Finally, we successfully applied BSA with cryopreserved progeny pools.

## 2. Methods

### 2.1. Ethics approval and consent to participate

The study was performed in strict accordance with the recommendations in the Guide for the Care and Use of Laboratory Animals of the National Institutes of Health (NIH), USA. To this end, the Seattle Children’s Research Institute (SCRI) has an Assurance from the Public Health Service (PHS) through the Office of Laboratory Animal Welfare (OLAW) for work approved by its Institutional Animal Care and Use Committee (IACUC). All of the work carried out in this study was specifically reviewed and approved by the SCRI IACUC.

### 2.2. Preparation of the genetic crosses

We generated the crosses using FRG NOD huHep mice with human chimeric livers and *A. stephensi* mosquitoes as described by Vaughan *et al.* ^22^. Two genetic crosses were generated for this study - Mal31×KH004 and NF54×NHP1337. Three individual recombinant pools were generated for each cross, by using different cages of infected mosquitoes. To start each cross, gametocytes from both parental parasite strains were diluted to 0.5% gametocytemia in a human serum erythrocyte mix, to generate infectious blood meals (IBMs). IBMs from each parent were mixed at equal proportions and fed to three cages of mosquitos (150 per cage).

We examined the mosquito infection rate and oocyst number per infected mosquito 7-10 days post-feeding. Fifteen mosquitoes were randomly picked from each cage and dissected under microscopy. Sporozoites were isolated from infected mosquito salivary glands and 2-4 million sporozoites from each cage of mosquitoes were injected into three FRG huHep mice (one cage per mouse), intravenously. To allow the liver stage-to-blood stage transition, mice are infused with human erythrocytes six and seven days after sporozoite injection. Four hours after the second infusion, the mice are euthanized and exsanguinated to isolate the circulating ring stage *P. falciparum*-infected human erythrocytes. The parasites from each mouse constitute the initial recombinant pools of recombinant progeny for genetic mapping experiments. We maintained the initial pools in AlbuMAX supplemented RPMI media; we genome sequenced aliquots from each pool to check allele frequencies from both parents. Two recombinant pools from each cross were used in this study for DHA BSA experiments.

### 2.3. Cryopreservation of progeny bulks

We expanded progeny bulks by growing in AlbuMAX supplemented RPMI for 7 days after the mouse bleeds. When cultures were 3% parasitemia and at least 70% ring-stage, progeny bulk stocks were cryopreserved in glycerolyte. Each 0.5ml cryopreserved stock contained ~100 μl packed red cells. We thawed and cultured parasites for 9 days prior to set-up in BSA experiments.

### 2.4. Measuring DHA resistance level using an eRRSA method

Cryopreserved stocks of the KH004 parent and the Mal31×KH004 uncloned bulk recombinant progeny pool and were thawed and grown in complete media (CM) [RPMI 1640 with L-glutamine (Gibco, Life Technologies), 50 mg/L hypoxanthine (Calbiochem, Sigma-Aldrich), 25 mM HEPES (Corning, VWR), 0.5% Albumax II (Gibco, Life Technologies), 10ug/ml gentamycin (Gibco, Life Technologies), 0.225% sodium bicarbonate (Corning, VWR)] at 5% hematocrit in O^+^ red blood cells (RBC) (Biochemed Services, Winchester, VA). Cultures were grown in separate flasks and maintained at constant pH, 7.0-7.5, temperature, 37°C, and atmosphere, 5% CO_2_/5% O_2_/90% N_2_. Cultures were kept below 2% parasitemia with media changes every 48 h. Parasites were synchronized to late stage schizonts using a single 70% Percoll density gradient as previously described ^23^. Four hours post-synchronization, parasitemia and stage were quantified using flow cytometry. 80 μl of culture and a 5% hematocrit RBC control were stained with SYBR Green I and SYTO 61 and measured on a Guava easyCyte HT (Luminex Co.). 50,000 events were recorded to determine relative parasitemia and stage. When at least 70% of parasites were in the ring-stage, the culture was diluted to 2% hematocrit and 0.5% parasitemia and parasites were aliquoted into a 96-well plate. Ten DHA concentrations were tested with a 2-fold dilution series starting with 2800 nM DHA. eRRSA assay set-up and qPCR amplification was done as previously described in ^23^ to calculate a fold change value between the untreated and treated sample for three biological replicates for each parasite.

### 2.5. DHA BSA and sample collection

Two recombinant pools from each cross (Mal31×KH004 or NF54×NHP1337) were selected and used for DHA BSA experiments. For each pool of recombinants, one sample was taken from the original progeny pool and was not synchronized (unsynchronized control), while three other samples were taken from the original progeny pool and synchronized at 0 h, 18 h, and 36 h (**Figure 3A**). We used a single 70% Percoll density gradient for synchronization (as described above) and allowed parasites to reinvade for four hours after synchronization to ring stages before determining parasitemia and stage via flow cytometry and setting up the DHA BSA experiment. All progeny pools (0 h, 18 h, and 36 h synchronized and unsynchronized control) were treated with DMSO (control), 50 nM DHA, or 100 nM and maintained in 48-well plates. After 6 hours, infected RBCs were washed three times with RPMI to remove any residual DHA or DMSO and then resuspended with new culture media to allow surviving parasites to grow out. Samples were collected right before drug treatment (day 0) and 5 days later when there was enough material for DNA extraction.

### 2.6. Library preparation and sequencing

We used Qiagen DNA mini kit to extract and purify the genomic DNA, and Quant-iT™ PicoGreen® Assay (Invitrogen) to quantify the amount of DNA.

For samples with less than 50ng DNA obtained, whole genome amplification (WGA) was performed before NGS library preparation. WGA reactions were performed following Nair et al ^24^. Each 25 μl reaction contained at least 5ng of *Plasmodium* DNA, 1× BSA (New England Biolabs), 1 mM dNTPs (New England Biolabs), 3.5 μM of Phi29 Random Hexamer Primer, 1× Phi29 reaction buffer (New England Biolabs), and 15 units of Phi29 polymerase (New England Biolabs). We used a PCR machine (SimpliAmp, Applied Biosystems) programmed to run a “stepdown” protocol: 35 °C for 10 min, 34 °C for 10 min, 33 °C for 10 min, 32 °C for 10 min, 31 °C for 10 min, 30 °C for 6 h then heating at 65 °C for 10 min to inactivate the enzymes prior to cooling to 4 °C. Samples were cleaned with AMPure XP Beads (Beckman Coulter) at a 1:1 ratio. We constructed next generation sequencing (NGS) libraries using 50-100 ng DNA or WGA product following the KAPA HyperPlus Kit protocol with 3-cycle of PCR. All libraries were sequenced at 150bp pair-end using Illumina Novaseq S4 or Hiseq X sequencers. We sequenced all bulk samples to a minimum coverage of 100×.

### 2.7. Mapping and genotyping

We individually mapped whole-genome sequencing reads for each library against the *P. falciparum* 3D7 reference genome (PlasmoDB, release32) using the alignment algorithm BWA mem (http://bio-bwa.sourceforge.net/) under the default parameters. The resulting alignments were then converted to SAM format, sorted to BAM format, and deduplicated using picard tools v2.0.1 (http://broadinstitute.github.io/picard/). We used Genome Analysis Toolkit GATK v3.7 (https://software.broadinstitute.org/gatk/) to recalibrate the base quality score based on a set of verified known variants ^25^.

After alignment, we excluded the highly variable genome regions (subtelomeric repeats, hypervariable regions and centromeres) and only performed genotype calling in the 21 Mb core genome (defined in ^22^). We called variants for each sample using HaplotypeCaller, and calls from every 100 samples were merged using CombineGVCFs with default parameters. Variants were further called at all sample-level using GenotypeGVCFs, with parameters: --max_alternate_alleles 6 --variant_index_type LINEAR --variant_index_parameter 128000 --sample_ploidy 2 -nt 20. We further filtered the variants calls by calculating the recalibrated variant quality scores (VQSR) of genotypes from parental parasites. Loci with VQSR less than 1 or not distinguishable between two parents were removed from further analysis.

The variants in VCF format were annotated for predicted functional effect on genes and proteins using snpEff v4.3 (https://pcingola.github.io/SnpEff/) with 3D7 (PlasmoDB, release32) as the reference.

### 2.8. Bulk segregant analysis

Only single-nucleotide polymorphism (SNP) loci with coverage > 30× that differed between the two parents were used for bulk segregant analysis. We counted reads with genotypes of each parent and calculated allele frequencies at each variable locus. Allele frequencies of one of the parents (NF54 or Mal31 in this study) were plotted across the genome, and outliers were removed following Hampel’s rule ^26^ with a window size of 100 loci. We performed the BSA analyses using the R package QTLseqr ^27^. Extreme-QTLs were defined when FDRs (Benjamini-Hochberg adjusted p-values) were less than 0.01 ^28^. Once a QTL was detected, we calculated and approximate 95% confidence interval using Li’s method ^29^ to localize causative genes.

## 3. Results

### 3.1. Genetic crosses

We employed *Anopheles stephensi* mosquitoes and human liver-chimeric FRG huHep mice as described in Vaughan *et al* ^22^, to generate two unique genetic crosses (Mal31×KH004 and NF54×NHP1337) for this study (**Figure 1A, Table 1**).

**Table 1.** Segregating variation in two genetic crosses. In addition to a known ART-R associated mutation in *kelch13*, four variants (underlined amino acid changes) showing strong associations in a genome wide association analysis ^6^ are also segregating in these crosses.

| Gene name | Gene ID | Cross 1 |  | Cross 2 |  |
| --- | --- | --- | --- | --- | --- |
|  |  | Cambodia KH004 | Malawi Mal31 | Thailand NHP1337 | Africa NF54 |
| *kelch13* | PF3D7_13437000 | C580Y | WT | C580Y | WT |
| *fd* | PF3D7_1318100 | <u>D193Y</u> | WT | <u>D193Y</u> | WT |
| *arps10* | PF3D7_1460900 | <u>V127M</u>,D128H | WT | <u>V127M</u>,D128H | WT |
| *mdr2* | PF3D7_1447900 | <u>T484I</u>,F423Y,G299D,S208N | I492V,S208N | <u>T484I</u>,F423Y,G299D,S208N | WT |
| *crt* | PF3D7_0709000 | DD2<sup>1</sup>+G367C | WT | DD2<sup>1</sup> | WT |

1. Alleles carry same amino acid mutations as the Dd2 parasite line (M74I, N75E, K76T, A220S, Q271E, N326S, I356T, R371I) when compared to reference (3D7).

#### Mal31×KH004

Mal31 is a *kelch13* wild-type parasite isolated from a patient in Malawi in 2016, KH004 is a *kelch13* mutant parasite carrying the common C580Y mutation and was isolated from Western Cambodia in 2016. There are 14,455 core-genome SNPs (defined in ^25^) distinguishing these two parental parasites. The gametocytes from both parental parasites were mixed in a ~50:50 ratio to infect approximately 450 mosquitoes. We generated four independent recombinant pools from this cross by using independent groups of 100 mosquitoes for the isolation of sporozoites and four individual FRG huHep mice that were infected with the sporozoites. Recombinants are generated after gametes fuse to form zygotes in the mosquito midgut (**Figure 1A**). Mitotic division of the four meiotic products ultimately leads to the generation of thousands of haploid sporozoites within each oocyst. The oocyst prevalence from the cross ranged from 71-91%, with an average of 10 oocysts per mosquito. The estimated number of recombinants for each pool were 4,000 (10 oocysts × 4 recombinants × 100 mosquitoes). The initial allele frequencies of Mal31 were 0.75-0.79 for bulk pools directly from mice; the deviation from the expected even representation of the two parental alleles indicates the existence of Mal31 selfed progeny generated by fusion of male and female gametes of Mal31. Thus, the expected unique recombinants per pool will be ≤ 4,000 for this cross. Two recombinant progeny pools were randomly selected for use in this study. For these two pools, the genome-wide allele frequencies shifted to 0.53 and 0.55 respectively after 15 days of *in vitro* culture prior to DHA BSA experiments (**Figure S1**). The changes to a more even representation of the two parental genomes suggests selection against selfed progeny during *in vitro* blood culture, which we have detected previously in a different cross ^15^.

#### NF54×NHP1337

NF54 is parasite of African origin, and NHP1337 is a C580Y *kelch13* mutant parasite isolated from the Thai-Myanmar border in 2011 and has been used previously in a genetic cross ^30^. There are 17,318 SNPs distinguishing the two parental parasites. The gametocytes from both parental parasites were mixed in a ~50:50 ratio to feed about 450 mosquitoes (150 per cage, three cages). We generated two independent recombinant pools for this cross by using different sets of mosquitoes. The oocyst prevalence for this cross was 90% (range: 80-100%), with an average burden of 25 oocysts per mosquito midgut (range: 14-40). The estimated number of recombinants for each pool used were 10,000 (25 oocysts × 4 recombinants × 100 mosquitoes). The allele frequencies of NF54 were 0.41 and 0.46 for bulk pools before DHA BSA experiments (**Figure S2**).

### 3.2. Determining DHA concentrations for BSA experiments

Maximal inhibitory concentrations, such as IC_50_ and IC_90_ would typically be used to determine the concentration of drugs used for BSA experiments (**Figure 1B**). However, for DHA, IC_50_ and IC_90_ values are not correlated with patient clearance half-life, the key clinical resistance readout ^31^. In this study, we used an assay we developed, eRRSA, to measure the DHA resistance level of the resistant parent KH004 and one of our recombinant progeny pools from the Mal31 ×KH004 cross. eRRSA relies on parasite growth fold change [2^(average *ct* drug treated – average *ct* control)^] to quantify parasite resistance to DHA and this read out is strongly correlated to patient clearance half-life ^23^. We generated DHA eRRSA dose-response curves, i.e., one eRRSA for each of 10 different drug concentrations (**Figure 2**) to identify concentrations of DHA that would kill some but not all of the sensitive parasites. The eRRSA dose response curves for resistant and pooled progeny were not significantly different (F_4,68_ = 1.518, *p* = 0.207). We chose 50 nM and 100 nM for our DHA BSA experiment: 50 nM and 100 nM correspond to an eRRSA fold change of 14 (46.1% parasite survival for KH004 and 52.7% parasite survival for the Mal31 × KH004 recombinant pool) and a fold change of 20 (32.4% parasite survival for KH004 and 34.1% parasite survival for the Mal31 × KH004 recombinant pool), respectively.

**Figure 2.**
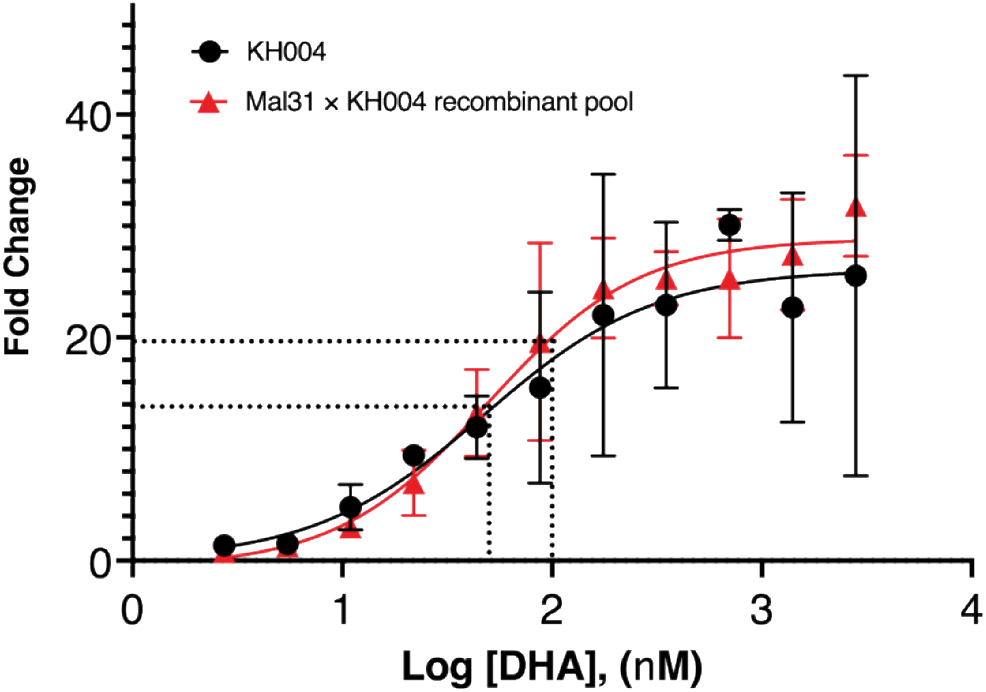
eRRSA dose-response curves for the DHA resistant parent KH004 and a Mal31 × KH004 recombinant progeny pool. Synchronized ring-stage parasites were treated with 10 DHA concentrations (in a two-fold dilution series starting at 2,800 nM) for 6 hours. Higher fold-change indicates increased parasite death in drug treated versus untreated parasites using our eRRSA method. Both the KH004 parent (artemisinin-resistant) and the Mal31×KH004 recombinant progeny pool were tested using three biological replicates to determine the optimal DHA concentration for bulk segregant DHA selection. Dashed lines indicate 50 nM and 100 nM were chosen as discerning doses for treatment of bulk populations to selectively enrich for resistant parasites.

### 3.3. BSA with pooled progeny from the Mal31×KH004 cross

Malaria parasites show different tolerance levels to artemisinin drugs at different lifecycle stages (ring, trophozoite or schizont) ^32^. To remove the influence of parasites from different stages and maximize the power of DHA BSA, we synchronized samples from the recombinant progeny pools. This was done at three time points (0, 18 or 36 h) prior to DHA BSA, and we also included an unsynchronized control (**Figure 3A**). High depth Illumina sequence (>100×) profiles of these four progeny pools revealed no significant differences (**Figure S3**).

**Figure 3.**
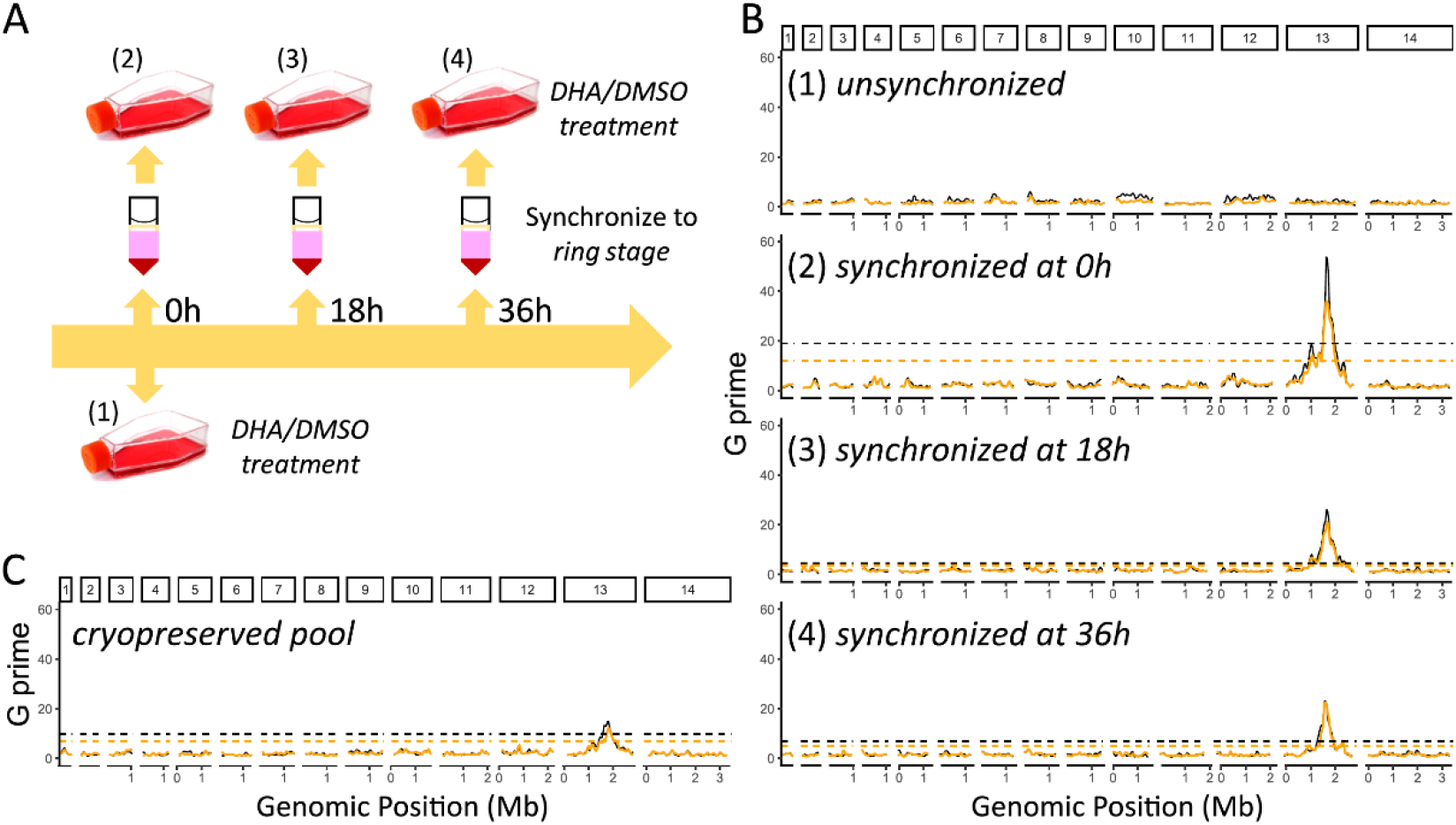
DHA bulk segregant analysis (BSA) on bulk progeny from the Mal31×KH001 cross. (A) Synchronization of parasites at the ring stage: (1) pools without synchronization; (2-4) pools synchronized at 0h, 18h and 36h, respectively. (B) DHA BSA analyses of pools synchronized at the different time points outlined in (A). (C) DHA BSA with a cryopreserved recombinant pool synchronized at 0h. Orange and black lines are G prime values comparing allele frequency of 50 nM or 100nM DHA treated pools with control pools; orange (50nM) and black (100nM) dashed lines are corresponding significance thresholds (FDR = 0.01).

We compared the Mal31 allele frequency from segregant pools with and without DHA treatment using genome-wide G’ ^28^ to measure the significance level of detected QTLs (**Figure 1 C&D**). For each detected QTL (FDR < 0.01), we calculated 95% confidence intervals to define the QTL boundaries (**Table S1**). For pools without synchronization, we were not able to detect any significant QTL. For pools that were synchronized, we identified a single strong QTL on chromosome 13 at the *kelch13* locus (**Figure 3, Figure S4**). The largest allele frequency difference between control and DHA treated parasites was detected from pools synchronized at 0 h. We did not detect significant differences between 50nM and 100nM DHA treatments.

### 3.4. BSA with cryopreserved pooled progeny from the Mal31×KH004 cross

Generation of recombinant progeny pools for each BSA experiment requires *in vitro* culture of parasites, maintaining *Anopheline* mosquitoes, and infection of FRG huHep mice, and is extremely labor intensive and costly. An alternative strategy is to use cryopreserved progeny pools for BSA experiments. We compared allele frequencies of a Mal31×KH004 progeny pool before and after cryopreservation. In the absence of drug treatment, the allele frequency across the genome remained at the same level in progeny pools before and after synchronization. We observed a modest divergence in allele frequency at chromosome 2 and 14, with increased frequency of Mal31 alleles, which may indicate a reduction in parasite population diversity (**Figure S3 B**). We applied DHA BSA to this cryopreserved pool (**Figure 3C**) and compared results with BSA before cryopreservation (**Figure 3B, panel 2**). We detected a strong QTL at the *kelch13* locus, although the G prime values were approximately five-fold less and the 95% CIs were approximately ten times larger as compared to the results from the original BSA analysis.

### 3.5. BSA with NF54×NHP1337 cross

Based on previously published eRRSA values for NHP1337 ^23^, we also applied DHA BSA to pooled progeny from the NF54×NHP1337 cross using the same methodology and DHA concentrations as described for the Mal31×KH004 cross. As with the Mal31×KH004 cross, no significant difference in allele frequency was detected from pools with and without synchronization prior to BSA (**Figure S2**). Similarly, we detected a strong QTL at the *kelch13* locus from both biological replicates after synchronization, but not in the unsynchronized control (**Figure S5**). A second smaller QTL peak on chr 14 was detected in one of the two biological replicates.

## 4. Discussion

Given the success of BSA approaches for identification of drug resistance loci in rodent malaria ^33–35^, our aim was to develop a BSA method to identify drug resistance loci in *P. falciparum* to better exploit our capacity to conduct *P. falciparum* genetic crosses in humanized mice ^36^. To develop our approach, we chose DHA, the active metabolite of artemisinin, as a test drug. This drug shows highly stage specific drug action, targeting young rings ^37^. Furthermore, there is a well-known locus (*kelch13*) underlying ART-resistance that provides a valuable positive control for assessing and optimizing our methods ^17^. We specifically aimed to explore the impact of (i) parasite synchronization, (ii) parasite cryopreservation, and (iii) to develop strategies to choose optimal drug selection regimens for BSA. The power of BSA is strongly dependent on the number of recombinant parasites present in progeny pools. Therefore, methods for BSA with malaria parasites should aim to minimize time in culture prior to selection, and avoid possible procedures that may reduce the diversity of recombinant pools.

### 4.1. Synchronization influences BSA success

Antimalarial drugs show a spectrum of specificity for different parasite asexual blood stages (rings, trophozoites, schizonts). At one end of the spectrum, drugs such as chloroquine kill all of these stages and show minimal stage specificity. At the other extreme, drugs such as DHA, cycloheximide, and trichostatin ^38^, or newly developed drugs like BCH070^39^, show strong specificity for early rings, while drugs such as lumefantrine (peak activity: 16-40h), mefloquine (16-40h) and piperaquine (12-36 h) do not kill early rings and show specificity for trophozoites ^40^. Importantly, we determined that synchronization is essential for a successful DHA BSA: with synchronization, we were able to clearly detect the known *kelch13* peak and without synchronization we detected no peaks (Figure 3B). Synchronizing parasites to early ring-stage allowed us to specifically target ring-stage resistance; the presence of other parasite stages in the culture is an additional variable that thus limits the success of selection. We suggest that synchronization should be used for BSA experiments with drugs like DHA that are active against early rings, and for drugs where no stage specificity information is available.

Our experiments with Mal31×KH004 also involved differently synchronized parasite pools (0, 18 or 36 h) prior to BSA. Selection on all three pools resulted in a strong significant QTL peak at *kelch13*. Notably, the strongest QTL localizing to *kelch13* was in the population synchronized at 0h and reduced in magnitude at 18h and 36h. This is likely due to the developmental stage differences between the recombinant progeny pools corresponding to these timepoints. According to flow cytometry of the progeny pool synchronized at 0h, the population was 91% ring stage; at 18h 89% of the synchronized population was ring stage, and at 36h 65% of the synchronized population was ring stage, and the unsynchronized population was 68% ring stage. We suspect that the high proportion of ring-stages in the 0 h synchronized progeny pool maintained higher diversity than the 18h and 36h synchronizations, which resulted in a stronger QTL. The lack of a detectable QTL in unsynchronized cultures, most likely results from presence of late stage parasites (late trophozoites and schizonts) which are not killed by DHA, but contain more DNA than rings and therefore swamp out signal from rings in the BSA experiments.

### 4.2. Cryopreserved recombinant progeny pools can be used for drug BSA experiments

*P. falciparum* asexual stage culture can be maintained and manipulated indefinitely *in vitro* or cryopreserved for future study. Once bulk recombinant progeny pools are isolated from the FRG huHep mouse, they can be directly used in BSA experiments, and they also can be cryopreserved. Cryopreservation allows for multiple BSA experiments to be conducted using the same recombinant progeny pool at a later date. However, only ring stage parasites survive the freeze and thaw process. Due to the asynchrony over time of the clones in the bulk recombinant progeny pools recovered from FRG huHep mice, the diversity of the population is likely to be diminished during the process of cryopreservation, thawing, and parasite regrowth. Encouragingly, these BSA experiments reveal the same QTL before and after cryopreservation (Figure 3C), confirming that sufficient genetic variation is maintained in cryopreserved stocks of recombinant progeny pools to be used for BSA experiments. However, we note that the chromosome 13 QTL peak detected, while strongly significant, is smaller than detected prior to cryopreservation, suggesting some loss of recombinants within pools. Methods that maximize size and diversity of progeny pools can increase the success of BSA experiments. These include ensuring that bulk progeny are cryopreserved when rings predominate, and that culture time between parasite recovery from cryopreservation and BSA is minimized. We also note that we are able to conduct BSA experiments using cryopreserved progeny pools from several biological replicates of each cross, providing replication and added confidence in the QTL peaks detected.

### 4.3. Determining drug concentrations for BSA selection experiments

The goal of a selective drug BSA is to kill only the sensitive parasites in the sample exposed to drug. Therefore, the drug concentration chosen for selection is very important. If the dose is too high, few parasites in the treated pool will survive, while if the dose is too low many sensitive parasites will survive, and one will not see enrichment of alleles and QTLs resulting from drug selection. In yeast BSA experiments, particular strong selection (recovers <1% of the total progeny) has been used as a rule of thumb to select progeny showing extreme phenotypes ^7^. This works well for yeast, where recombinant pools can contain >10^6^ recombinants, but in our experience with *P. falciparum,* the use of such stringent drug doses for BSA can result in poor recovery of parasites in the treated pools.

To choose drug concentrations for BSA, we measured drug response in the resistant parent and one of our recombinant progeny pools. Traditional IC_50_ measures work poorly with short half-life drugs such as DHA, so we examined dose response using a modified ring stage assay (eRRSA; Figure 2) ^23^ which examines parasite survival following a short pulse of drug exposure, to generate dose response curves. We observed a slight shift in the eRRSA dose-response curve between the resistant parasite, KH004, and the Mal31 × KH004 recombinant progeny pool, although the two curves were not significantly different. We chose two DHA doses at 50 nM and 100 nM for our DHA BSA experiment to kill some but not all the sensitive parasites. Given the high value of stock samples of bulk progeny and the time and cost of the BSA experiment, it is important to conduct these initial experiments to enhance the chance of successful selections. Encouragingly, we found that both DHA doses revealed the same QTLs with near equivalent results with this small dosing concentration range of DHA in the Mal31×KH004 cross. The 100nM dose (32.4% parasite survival for KH004 and 34.1% parasite survival for the Mal31 × KH004 recombinant pool) revealed slightly higher QTL peaks and having two doses provided technical replication. Use of dose response data from uncloned progeny pools could be helpful when progeny drug responses fall outside the range of the parents (transgressive segregation)^41^ or in cases where there is no prior information about progeny drug responses. A practical alternative that should work when the genetic architecture is simple and parent phenotypes are significantly different is to measure drug responses in parental parasites, and choose two or three drug doses bracketing the IC_50_ (of equivalent metric) of the most resistant parent. We anticipate that a use of a well-chosen range of selection doses could reveal aspects of the genetic architecture that would not be revealed by a single dose.

### 4.4. Genetic architecture of DHA resistance

The primary goal of this project was to develop methodology for drug BSA experiments with *P. falciparum*. However, these crosses also provide information on the genetic architecture of artemisinin resistance. Four lines of evidence suggest that loci other than *kelch13* may contribute to resistance: (i) GWAS studies have identified several loci (*ferredoxin, apicoplast ribosomal protein S10, multidrug resistance protein 2* and *crt*) as well as *kelch13* to be associated with artemisinin resistance ^6^; (ii) parasites showing slow clearance from artemisinin-treated patients but lacking mutations in *kelch13* have been described ^42^; and (iii) longitudinal analyses have shown that other loci show parallel changes in allele frequency over time ^43^, suggesting that other loci may also contribute to slow clearance phenotypes; (iv) *in vitro* selection has revealed other loci, such as *pfcoronin,* that can modulate ART-R ^21^.

The two crosses analyzed here show a single QTL indicating that *kelch13* is the largest genetic signal of this phenotype. We observed smaller signals meeting significance threshold on chromosomes 14 for some replicates (Fig .S5). However, as these reach significance in only some replicates, and are also seen in control (untreated samples), they most likely reflect loci underlying differences in growth rate, rather than try resistance QTLs. This illustrates a further advantage of BSA: by conducting replicate BSA experiments using independent pools of progeny we can minimize detection of false positive QTLs.

Importantly, these crosses allow us to interrogate candidate loci. Both genetic crosses analyzed have segregating SNPs in *ferredoxin, apicoplast ribosomal protein S10, multidrug resistance protein 2* and *crt.* Interestingly, while these loci show strong association with slow clearance in a GWAS, they were proposed to provide permissive background mutations on which ART-R mutations could arise ^6^, perhaps through impacts on fitness. In our BSA experiments, we sampled parasite pools at several time points after drug exposure. This allows detection of QTLs underlying both drug resistance and fitness. We did not detect QTLs at any of these four loci, suggesting that these SNPs do not play a role in ART-R, or contribute to fitness of ART-R parasites. These results are consistent with results from CRISPR/Cas9 editing of *ferrodoxin* and *mdr2* SNPs ^44^, where no impact of these SNPs on ART-R or fitness was found.

Undertaking more complex BSA analyses using an increased range of drug concentrations may help uncover further loci that play a role in resistance. In addition, further crosses may identify additional loci underlying artemisinin resistance. For example, for 20 years *crt* has been considered to be the primary determinant of CQ resistance. However, in parallel work on CQ-resistance we have identified a prominent chromosome 6 QTL that acts in concert with *crt* to determine CQ resistance in some crosses ^45^. Our ability to conduct genetic crosses in FRG huHep mice provides a direct approach to identify suspected *non-kelch13* mechanisms of artemisinin resistance using parasites isolated from patients ^46^.

### 4.5. BSA is broadly applicable to selectable phenotypic traits

We have demonstrated that BSA is a powerful approach in determining genes related to drug resistance (this study) in *P. falciparum*, paralleling findings from other organisms ^47–50^. In other work we have used BSA to examine the impact of culture conditions (serum or AlbuMAX) ^16^, and identifying genome regions that impact parasite fitness ^15^. We anticipate that BSA can also be explored to study any phenotypic trait that can be measured by selection. For example, we can examine the selection on parasite populations at different temperatures, which simulate one of the malaria symptoms – fever ^51^. We can control the pH of culture media, which can be changed in patients with severe malaria due to acidosis ^52^. We can also study the selection of human immunity on progeny population by culturing parasites with serum from recovered patients or people that have never had malaria, as was done with mouse malaria ^53^. BSA can also be used to study parasite-host interactions. Malaria parasites invade human red blood cells (RBCs) by a series of interactions between host and parasite surface proteins ^54^. Hence, we can use different blood cell populations to examine genes underlying parasite invasion phenotypes. Furthermore, Anopheline mosquitoes are the intermediate host between parasite and human and the distribution of mosquitoes varies at different geographical locations. For example, *Anopheles gambiae* mosquitoes are common vectors in Africa, while *Anopheles dirus and Anopheles minimus* are the dominant vectors in Asia ^55^. Hence we can infect different mosquitoes with recombinant progeny populations to examine the selection at the mosquito stage, as pioneered by Molina-Cruz et al ^56^.

### 4.6. Technical considerations & caveats

Statistical power and numbers of recombinants. The power of BSA is limited by the number of unique recombinant progeny in a pool. To maximize the number of recombinants, we can increase the number of mosquitoes we use for sporozoite isolation per cross, thereby increasing the number of unique recombinants. In addition, we are not limited to one mouse per cross and then can thus increase mouse numbers and infect each mouse with unique sporozoite pools, providing biological replicates. Cryopreservation of large volumes of recombinant after the transition to *in vitro* culture is also important and upon thawing, multiple independent pools can be analyzed with downstream BSA. Selfed progeny are another limitation for the statistical power of BSA. We have found large numbers of inbred progeny generated by mating between male and female gametes from the same parent, in some crosses ^15,57^. However, selfed progeny are selectively removed during asexual growth ^15^. Selfed progeny could also be removed by carrying out crosses with parental strains that only produce either functional male or female gametes, to ensure that all progeny are recombinant. For example, treatment with aphidicolin blocks male game production ^58^.

Assessing the numbers of recombinants. We have been using the number of oocysts and sequences from cloned progeny to estimate the maximum number of recombinants in a recombinant pool. Measuring the number of recombinant progeny and the proportion of inbreeding in a pool before BSA could be particularly useful for future crosses. This can be done by laborious cloning from the same pools ^30^. Alternatively, single cell genome sequencing ^59^ can be employed to analyze the diversity of the recombinant pool. Thus, rapid genotyping of sporozoites or the progeny pool could provide details on the dynamics of BSA experiments, which will then allow for the further optimization approaches ^60^.

Balancing the experiment time and culture volume. BSA can be used to examine the impact of multiple different selections in parallel. However, if multiple experiments are conducted at the same time, large numbers of parasites are needed, which requires a longer culture time. Population diversity decreases during *in vitro* culture due to parasite competition ^15^. A solution to this is to use smaller culture volumes for BSA experiments without compromising genetic diversity. We have used 3 ml (6-well plate, in ^15,16^) and 500 μl culture (48-well plate, in this study) at 1% parasitemia for our previous experiments, which both allow us to collect enough material for sequencing and keep diversity of the bulk. However, we were not able to sample efficiently using smaller culture volumes such as 100 μl from 96-well plate.

## 5. Data Availability

All data needed to evaluate the conclusions in the paper are present in the paper and/or the Supplementary Materials. All raw sequencing data have been submitted to the NABI Sequence Read Archive (SRA, https://www.ncbi.nlm.nih.gov/sra) under the project number of PRJNA524855. Additional data related to this paper may be requested from the authors.

## 6. Acknowledgments

This work was supported by National Institutes of Health (NIH) program project grant P01 AI127338 (to MF) and by NIH grant R37 AI048071 (to TJCA). Work at Texas Biomedical Research Institute was conducted in facilities constructed with support from Research Facilities Improvement Program grant C06 RR013556 from the National Center for Research Resources. SMRU is part of the Mahidol Oxford University Research Unit supported by the Wellcome Trust of Great Britain. The parental line, Mal31, used in the *Mal31×KH004* cross was sampled from a Malawian patient in 2016 as part of a cross-sectional study funded by the Wellcome Trust of Great Britain (Grant no. 099992/Z/12/Z to SCN). We thank the patients who provided parasites used in this work.

## Declaration of interests

We declare no competing interests.

## Contributors

TJCA, MTF and AMV conceived the project. KVB, XL and SK conceptualized and planned the study. SN, FN, MD, RT, TP, DL provided and characterized malaria parasites from infected patients from which parental parasites were chosen. SK, BAA and MTH prepared all the genetic crosses. KVB, LAC and DAS prepared BSA samples and DHA drug assay experiment. XL interpreted and visualized the data and performed formal analysis. ED and AR were involved in next generation sequencing library preparations. KVB, XL, SK, AMV, MTF and TJCA wrote the original manuscript. All authors read and approved the final manuscript.

## Notes

### Competing Interest Statement

The authors have declared no competing interest.

