## Supplementary material for "Optimizing bulk segregant analysis of drug resistance using *Plasmodium falciparum* genetic crosses conducted in humanized mice": bsa.optimization.11.24.2021_bioRxiv_SI.pdf

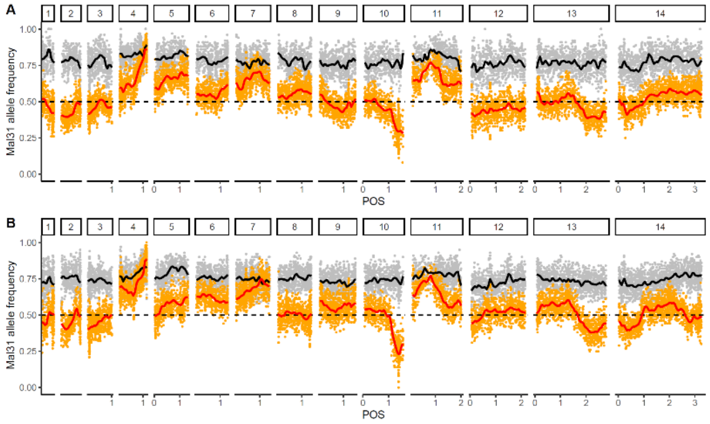


**Figure S1**. Genome-wide allele frequency of Mal31 in the Mal31×KH004 crosses in initial recombinant pools (grey dots and black lines), and pools after 15 days of *in vitro* culture that were used for CQ BSA experiments (orange dots and red lines). Plots in A and B panels showed allele frequency from different recombinant pools (biological replicates).

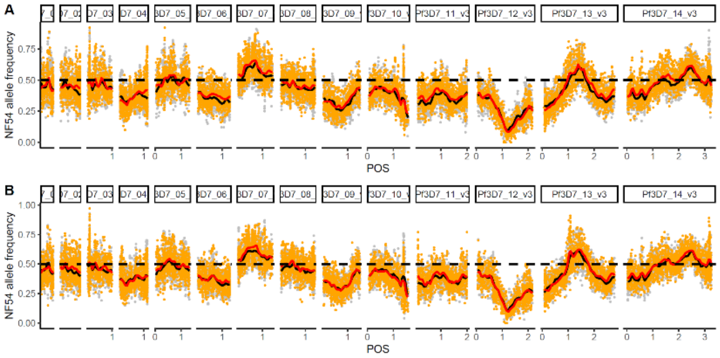


**Figure S2**. Genome-wide allele frequency of NF54 in the NF54×NHP1337 crosses in initial recombinant pools (grey dots and black lines), and pools after synchronization (orange dots and red lines) that were used for BSA experiments. Plots in A and B panels showed allele frequency from different recombinant pools (biological replicates).

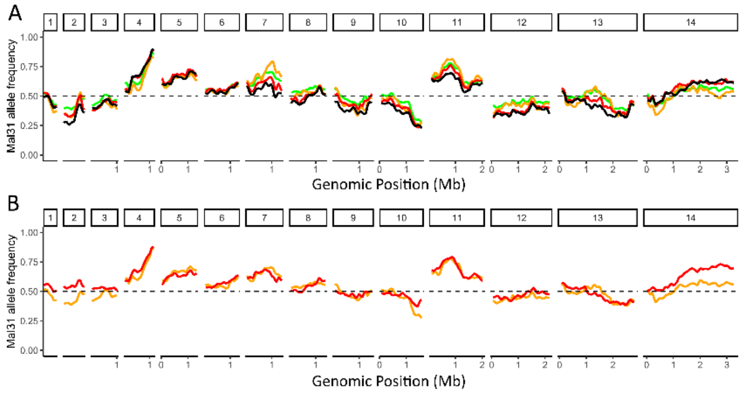


**Figure S3**. Genome-wide allele frequency of Mal31 before/after (A) synchronization and (B) cryopreservation. A, black lines indicate Mal31 allele frequency before synchronization; green, orange and red lines are Mal31 allele frequency after synchronization at 0h, 18h and 36h (see Figure 3 for details). B, Mal31 allele frequency before (orange) and after (red) cryopreservation.

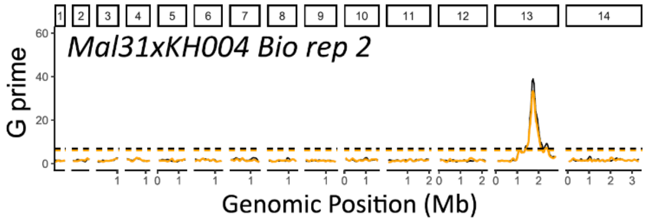


**Figure S4**. DHA bulk segregant analysis with cross Mal31xKH004 biological replicate 2. Recombinant pool was synchronized at 0h. Red and black lines are G prime values comparing allele frequency of 50nM or 100nM DHA treated pools with control pools; dashed lines are corresponding significance thresholds (FDR = 0.01).

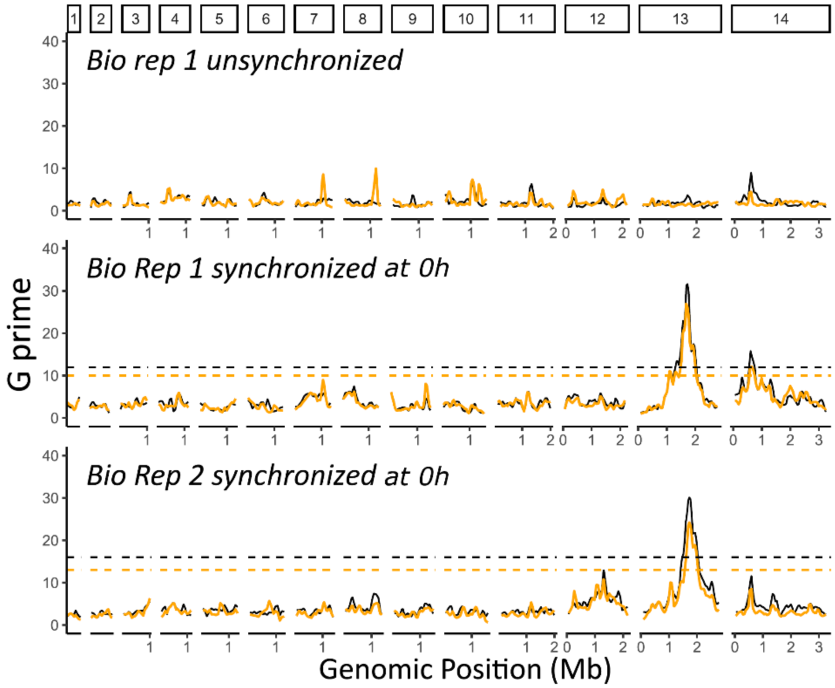


**Figure S5**. DHA bulk segregant analysis with cross NF54xNHP1337. Red and black lines are G prime values comparing allele frequency of 50nM or 100nM DHA treated pools with control pools; dashed lines are corresponding significance thresholds (FDR = 0.01).

**Table S1.** 95% confidence interval (CI) from different BSA analysis.

| Cross | BSA | Lower CI | Upper CI | Peak | CI size (kb) |
|---|---|---|---|---|---|
| Mal31xKH004 | M3.50nM.P0 | 1,646,404 | 1,803,513 | 1,718,368 | 157.109 |
| Mal31xKH004 | M3.100nM.P0 | 1,663,525 | 1,760,723 | 1,697,887 | 97.198 |
| Mal31xKH004 | M3.50nM.P18 | 1,559,170 | 1,803,513 | 1,723,034 | 244.343 |
| Mal31xKH004 | M3.100nM.P18 | 1,618,296 | 1,793,121 | 1,697,887 | 174.825 |
| Mal31xKH004 | M3.50nM.P36 | 1,618,296 | 1,813,639 | 1,706,634 | 195.343 |
| Mal31xKH004 | M3.100nM.P36 | 1,643,763 | 1,793,121 | 1,697,887 | 149.358 |
| Mal31xKH004 | M3.50nM.P0.frozen | 896,459 | 2,569,833 | 1,766,359 | 1673.374 |
| Mal31xKH004 | M3.100nM.P0.frozen | 1,119,676 | 1,971,214 | 1,766,359 | 851.538 |
| Mal31xKH004 | M4.50nM.P0 | 1,663,525 | 1,776,808 | 1,723,034 | 113.283 |
| Mal31xKH004 | M4.100nM.P0 | 1,663,525 | 1,773,273 | 1,723,034 | 109.748 |
| NF54xNHP1337 | M11.50nM.P0 | 1,596,756 | 1,792,824 | 1,681,767 | 196.068 |
| NF54xNHP1337 | M11.100nM.P0 | 1,661,966 | 1,792,824 | 1,724,204 | 130.858 |
| NF54xNHP1337 | M21.50nM.P0 | 1,618,296 | 1,912,767 | 1,724,204 | 294.471 |
| NF54xNHP1337 | M21.100nM.P0 | 1,618,296 | 1,820,881 | 1,725,259 | 202.585 |
